# Detailed splenic single-cell biodistribution of phosphatidylglycerol-containing liposomes

**DOI:** 10.1101/2025.09.15.676298

**Authors:** Deja Porenta, Estefanía Lozano-Andrés, Enrico Mastrobattista, Femke Broere, Naomi Benne

**Affiliations:** Utrecht Institute for Pharmaceutical Sciences, Department of Pharmaceutics, Faculty of Science, Utrecht University, The Netherlands; Department of Biomolecular Health Sciences, Infectious Diseases and Immunology Division, Faculty of Veterinary Medicine, Utrecht University, The Netherlands; Division Cell Biology, Metabolism & Cancer, Department of Biomolecular Health Sciences, Faculty of Veterinary Medicine, Utrecht University, Utrecht, the Netherlands

**Keywords:** Antigen-presenting cells, liposomes, biodistribution, flow cytometry, spleen, autoimmunity

## Abstract

Antigen-specific tolerance induction is a promising therapeutic strategy for autoimmune and chronic inflammatory diseases. This can be achieved by targeted activation of regulatory T and B cells via antigen-presenting cells (APCs) in a tolerogenic context. Anionic antigen-carrying liposomes have shown potential, however, their efficacy is highly dependent on the administration route and liposomal rigidity. Here, we investigate the biodistribution and splenic APC subset-specific uptake of rigid DSPC:DSPG:CHOL liposomes compared to flexible DOPC:DOPG:CHOL liposomes using high-parameter flow cytometry. We developed a panel enabling identification of rare splenic APC subsets involved in immune tolerance, including CD169^+^ and MARCO^+^ marginal zone macrophages, red pulp macrophages, and conventional/plasmacytoid dendritic cells. Our findings confirm that rigid liposomes predominantly accumulate in the liver and spleen following IV injection, with negligible uptake in lymph nodes or lungs. Importantly, systemic distribution is significantly inhibited by subcutaneous administration, which is essential for tolerance induction. Among splenic APCs, macrophage subsets are major contributors to liposome uptake, though the liver remains the primary site of accumulation and may play a more dominant role in tolerance induction. This study underscores the importance of both liposomal design and delivery route in optimizing nanoparticle-based immune modulation strategies.

## Introduction

Antigen-specific tolerance induction is one of the highest goals for the treatment of autoimmune diseases and may be achieved by instructing the immune system to generate antigen-specific regulatory T and B cells (Tregs and Bregs, respectively) (1). To do this, the antigen must be presented to these cells by antigen-presenting cells (APCs), such as dendritic cells (DCs) and macrophages, in the context of immune tolerance, i.e. immune-suppressing cytokines or molecules (2). A promising strategy for modulating immune responses is to use drug delivery vehicles such as liposomes for antigen and tolerance-inducing signals co-delivery directly to the APCs (3). Liposomes represent a promising vehicle for drug delivery due to their biocompatibility and low toxicity (4). We have previously shown that anionic distearoylphosphatidylglycerol (DSPG)-containing liposomes that carry a disease-specific antigen can prevent the progression of autoimmune diseases such as rheumatoid arthritis (5) and atherosclerosis (6) in mouse models after intravenous (IV) or intraperitoneal (IP) injection. Moreover, others have shown successful tolerance induction in a mouse model of diabetes using human insulin peptides-loaded phosphatidylserine (PS)-containing liposomes (7). We have, however, not been successful in inducing tolerance with DSPG-liposomes using alternative administration routes, such as subcutaneous (SC) and intramuscular (IM) injection (data not shown). Others have also observed this effect (8), which suggests that the liposomes must enter the circulation to exert their tolerogenic effects. Studies investigating the organ distribution of fluorescently labelled liposomes show that IV-injected liposomes are predominantly cleared by tissue-resident macrophages (9); Kupffer cells in the liver (10), and red pulp macrophages (RPMs) in the spleen (11). Upon SC injection, however, a large fraction of the injected dose remains at the site of injection (SOI), and is thus unavailable to liver and spleen APCs (12).

APC subsets in the spleen and liver have been recognized as targets for immunotherapies aiming at immune tolerance induction (reviewed in (3,13)). The spleen is a specialized immunological organ and an important site for the recognition and uptake of apoptotic cells, pathogens, and other blood-borne materials, and red blood cell recycling (14,15). As the blood reaches the spleen through the branching splenic artery, it enters the marginal zone (MZ) and the white pulp. The white pulp is a lymphoid tissue with B and T cell zones, and is a key site for the initiation of adaptive immune responses (15,16). The majority of the blood that leaves the MZ enters the red pulp surrounding the white pulp, where RPMs filter and remove dying blood cells and foreign antigens. The MZ – located at the interface between the two compartments – hosts diverse innate immune cells, strategically positioned for efficient uptake of antigens from the circulation (16–18). MZ-resident macrophage subsets are known to play a vital role in peripheral self-tolerance (19,20) and preventing autoimmunity in mice (21,22) and humans (23,24). A CD169-expressing subset of macrophages reportedly interacts with conventional DC type 1 (cDC1s) to instruct CD8^+^ T-cell responses (25–27), and macrophage receptor with collagenous structure (MARCO)-expressing macrophages capture antigens for presentation to MZ B cells as well as regulate MZ B cell trafficking and organization of B cell follicles in the spleen (28–30). Depletion of either MZ macrophage population accelerated the progression of autoimmune diseases in mice, indicating their relevance in efferocytosis and maintaining self-tolerance (31,32). Besides macrophages, splenic DCs are a heterogeneous group of professional APCs and represent an important target for immunotherapies (13,33). The major populations are cDC1s, cDC2s, and plasmacytoid DCs (pDCs). cDC1s are known for their cross-presenting capacity and close association with CD8^+^ T cells (34,35), while cDC2s appear to be more heterogeneous and mainly present exogenous antigens to CD4^+^ T cells and drive Th2 and Th17 immune responses (36,37). Lymphoid progenitor-derived pDCs are a unique and heterogeneous DC population with multiple functional roles, such as mediating self-tolerance as well as antiviral and anti-tumor immunity (38–42). Besides macrophages and DCs, B cells, neutrophils, eosinophils, monocytes, and other myeloid cells also possess antigen uptake and presentation ability. While B and T cells are abundant in the spleen (making up 40-50% and 30-50% of all CD45^+^ immune cells, respectively)(43), innate immune cells correspond to a low percentage of splenocytes, i.e., CD169^+^ macrophages (∼0.05%) (44,45), MARCO^+^ macrophages (∼1.5%) (46), RPMs (∼3-7%) (47–49), cDC1s (∼0.5%) (50,51), cDC2s (∼1%), pDCs (∼0.5%) (43), and neutrophils (∼1%) (43). Despite their low abundance, they appear to be indispensable in their functional roles.

Although organ distribution and clearance of circulating liposomes and other drug delivery vehicles have been assessed before (52–57), comprehensive single-cell subset-specific distribution of liposomes remains largely unexplored. Cell heterogeneity in complex tissues requires single-cell analysis techniques to elucidate possibly rare events of liposome uptake in low-abundance cell types. Different techniques can be used to study the cellular distribution of fluorescently-labeled particles and drugs, such as immunofluorescence, multiphoton microscopy, and flow cytometry (52,58–60). Flow cytometry allows high-throughput cell immunophenotyping and the detection of liposome fluorescence per single cell, including the fluorescence intensity, which can be translated to the relative abundance of liposomes per cell (**Figure 1**). Following a preliminary experiment assessing the whole tissue biodistribution of IV-injected DSPC:DSPG:CHOL liposomes in mice after 1 h, we found that the liposomes accumulate in the liver and spleen, with no fluorescence signal observed in the lymph nodes (popliteal, inguinal, mesenteric, and brachial) and lungs (data not shown). Therefore, we identified the liver and spleen as the main organs of liposome accumulation one hour after IV injection. To study biodistribution of liposomes across heterogeneous splenic APCs, we developed an accessible flow cytometry panel for the identification of splenic immune cells in mice, considering relevant flow cytometric studies up to date (48,61–65). Specifically, we focus on APCs likely involved in liposome uptake and tolerance induction, such as DC and macrophage subsets. Through selection of a critical number of phenotypic markers we identified nine surface markers that were sufficient for simultaneous characterization of several subsets of macrophages – CD169^+^ marginal zone macrophages, MARCO^+^ marginal zone macrophages, and red pulp macrophages (RPMs), subsets of DCs – plasmacytoid DCs (pDCs), conventional DCs – DCs type 1 (cDC1s), and cDC2s, and other immune cells such as neutrophils, B cells, T cells, and other myeloid/monocytic cells.

**Figure 1:**
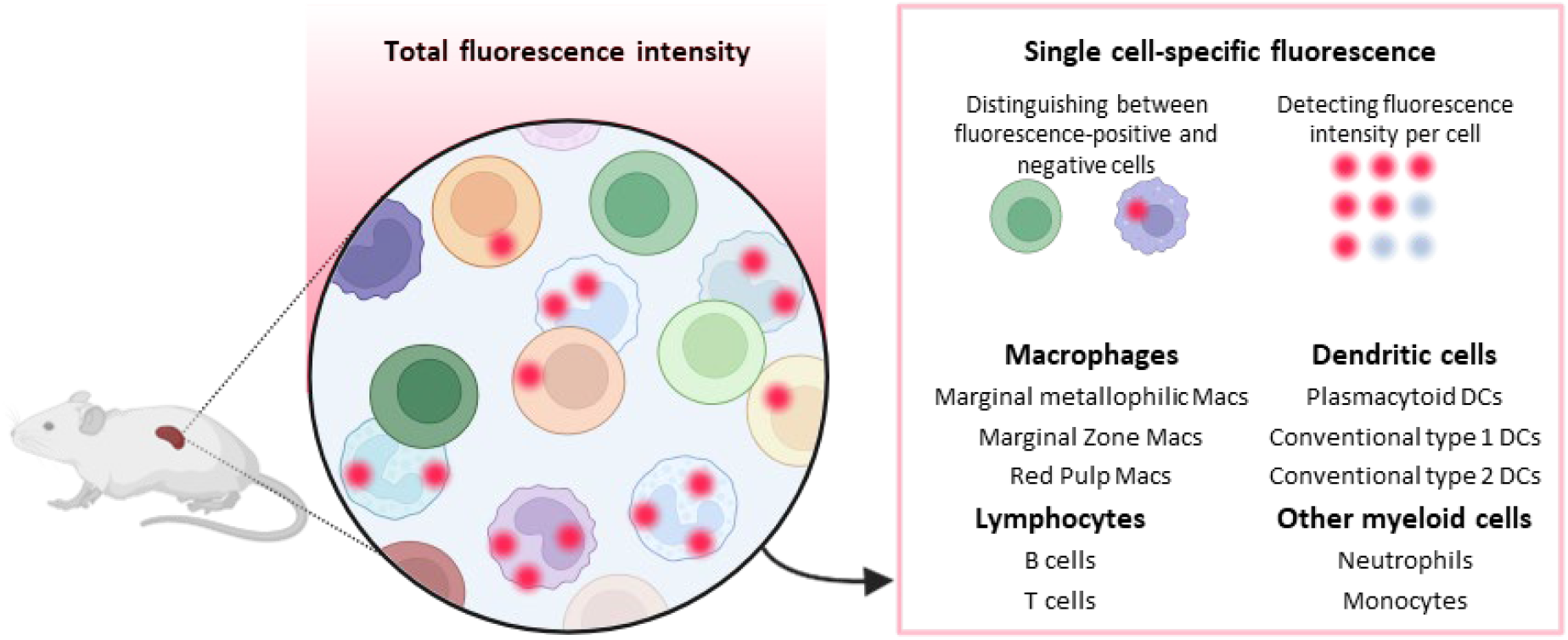
Fluorescent liposome biodistribution across splenic antigen-presenting cells, measured using single-cell flow cytometry. In contrast to whole-organ fluorescence, single-cell flow cytometry allows immunophenotyping of cell subsets as well as detecting fluorescence-positive cells and quantifying the relative amount of fluorescence per single cell.

Rigid DSPG-containing liposomes can induce tolerance; however, it remains unclear which splenic immune cells are involved in liposome uptake. Therefore, the goal of this work is to study the whole organ and splenic cell subset-specific distribution of rigid DSPC:DSPG:CHOL. Preliminary single-cell flow cytometry analysis allowed us and others (52,60) to detect splenic cell subsets contributing to the liposome uptake. To gain more insight into the processes that govern antigen-specific immune tolerance induction, different routes of administration (IV; 1h and 24h, SC; 24h) are compared. Besides the administration route, the physicochemical properties of liposomes and other nanoparticles, such as rigidity, size, and charge, have been shown to play an important role in the uptake and therapeutic efficacy (66–68). Liposomal rigidity largely depends on the phospholipid tail length and saturation (69,70). While incorporation of lipids with 18-carbon saturated tails (18:0), such as DSPC or DSPG yields a more rigid liposome, unsaturated phospholipids, e.g., DOPC or DOPG (18:1), form a more flexible liposome. While rigid DSPC:DSPG:CHOL liposomes can induce immune tolerance *in vivo*, this is not the case for DOPC:DOPG:CHOL liposomes, despite similar charge and size (69). Therefore, we additionally assessed the distribution of less rigid DOPC:DOPG:CHOL liposomes 1h after IV injection. Using flow cytometry, we investigated the involvement of specific splenic cell subsets, predominantly macrophage cell subsets, in the uptake of functionally tolerogenic anionic liposomes. Still, we observed that, at an organ level, the liver remains the major site of liposome accumulation.

## Methods

### Liposome formulation and characterization

Liposomes composed of DSPC (Avanti Polar Lipids, Birmingham, AL, USA), DSPG (Avanti Polar Lipids, Birmingham, AL, USA), and CHOL (Sigma-Aldrich, Burlington, MA, USA), or DOPC (Avanti Polar Lipids, Birmingham, AL, USA), DOPG (Avanti Polar Lipids, Birmingham, AL, USA), and CHOL were prepared via the dehydration/rehydration method. Briefly, 45 mg total of dry powder DSPC:DSPG:CHOL in a 4:1:2 molar ratio or DOPC:DOPG:CHOL in a 4:1:2 molar ratio was weighed and transferred to a sterile 50 mL round-bottom flask. 0.025 mol% DSPE-Cy5 (Avanti Polar Lipids) was added to each formulation for fluorescent labelling. Chloroform and methanol were added until the lipids were fully dissolved. The solvents were evaporated under vacuum in a rotary evaporator for 1 h at 40°C. The lipid film was placed under a N_2_ stream for 2h at RT to remove any residual solvents. 1 mL of phosphate buffer (10 mM, pH 7.4) was added to the lipid films and the suspension was homogenized in a water bath at 60°C for 1h. The liposomes were sized by high-pressure extrusion (LIPEX Extruder, Northern Lipids Inc., Burnaby, BC, Canada) at 60°C (DSPC:DSPG:CHOL:DSPE-Cy5 liposomes) or RT (DOPC:DOPG:CHOL:DSPE-Cy5 liposomes). The dispersion was passed four times through stacked 400 and 200 nm pore-size membranes (Whatman NucleoporeTM, GE Healthcare, Amersham, UK). The Z-average diameter and polydispersity index (PDI) of the liposomes were measured by dynamic light scattering (DLS) using a NanoZS Zetasizer (Malvern Ltd., Malvern, UK). The ζ-potential was measured by laser Doppler electrophoresis (Malvern Ltd.) using a universal dip cell. For both measurements, 10 µL of liposomes were diluted in 990 µL phosphate buffer. Samples were equilibrated to 25°C before measurement.

### In vivo biodistribution

8-week-old female WT mice on Balb/cAnNCrl background were purchased from Charles River Laboratories. Mice were randomized into experimental groups (N=3) and PBS group (N=4) based on weight. Mice were injected with 1.5 mg total lipid DSPC:DSPG:CHOL:DSPE-Cy5 liposomes (based on optimization), 1.5 mg total lipid DOPC:DOPG:CHOL:DSPE-Cy5 liposomes, an equivalent amount of free DSPE-Cy5 label or PBS. All groups were injected IV and an additional group was injected with DSPC:DSPG:CHOL:DSPE-Cy5 liposomes subcutaneously SC. Mice were sacrificed *via* cervical dislocation after 1 hour or 24 hours and perfused via the portal vein with 10 mL PBS. Animals were kept under standard housing conditions of the animal facility at Utrecht University, had access to a chow diet and water *ad libitum*, and all experiments were approved by the Animal Experiment Committee of Utrecht University (AVD10800202115687).

### Evaluation of whole organ fluorescence

One hour or 24 hours after injection, the lungs, liver, spleen, blood, femurs, tibia, popliteal lymph nodes (pLNs), inguinal LN (iLNs), brachial LN (bLNs) and mesenteric LN (mLNs) were collected. The whole spleen and liver was weighed, a small part of each was collected, weighed, and used for further processing. The bone marrow was isolated from the femurs and tibia of both hind legs and combined. All organs except liver were homogenized by passing through a 70 μm cell strainer (Corning, New York, USA). Erythrocytes were lysed in splenocyte samples by incubating the cell suspension in 2 mL Ammonium-Chloride-Potassium (ACK) lysis buffer (0.15 M NH4Cl, 1 mM KHCO3, 0.1 mM Na2EDTA; pH 7.3) for 1 minute. The liver was mechanically processed using blades, and digested in Collagenase D solution (Roche) (0,75 mg/ml) for 30 min at 37 °C to obtain a single-cell suspension. The cell suspension was centrifuged at 400x*g* for 5 min at 4 °C, and the resuspended pellet was passed through a 70 μm cell strainer (Corning, New York, USA). The cell suspension was centrifuged at 400x*g* for 5 min at 4 °C and erythrocytes were lysed by incubating the cell suspension in 2 mL Ammonium-Chloride-Potassium (ACK) lysis buffer (0.15 M NH4Cl, 1 mM KHCO3, 0.1 mM Na2EDTA; pH 7.3) for 1 minute. The suspension was centrifuged at 400x*g* for 5 min at 4 °C, and a part of the resuspended cells was collected for measurement. For LNs, the whole organ suspension was used. For the other organs and blood, a known amount of cells or volume was used, for later calculation to the whole organ. Blood volume was calculated based on each mouse’s weight, and bone marrow volume was assumed based on published data (71). All samples were added to clear-bottom, black-walled, 96-well plates (Greiner Bio-one, Solingen Germany). The Cy5 signal was measured using a microplate reader (Tecan Infinite M200 microplate, Tecan Trading, Männedorf, Switzerland) at RT at 630/670nm excitation/emission wavelength in bottom-reading mode using optimal gain values calculated based on the entire plate. The values per organ of PBS-injected mice were subtracted as background; for some organ samples this subtraction led to a negative value. The % dose was calculated based on the total fluorescence per organ/fluorescence of the injected dose. Non-recovered fluorescence was assumed to be cleared or in the case of SC injection at the site of injection (SOI).

### Spleen processing for flow cytometry

Part of the spleen was used for single-cell analysis of liposome association and uptake in APC subsets by flow cytometry. Splenocytes were processed as described above. The lysis was quenched by adding 9 mL of medium. The cells were centrifuged at 300x*g* for 5 min at 4°C, and the supernatant was removed. The pellet was resuspended in 5 mL of ice-cold medium. For counting, 10 μL of the cell suspension was diluted with 80 μL of medium and 10 μL of trypan blue, mixed, and 10 μL were transferred into a cell counting chamber. For flow cytometry staining, 0.6-1 million live splenocytes were seeded per well in a 96-well V-bottom plate.

### Flow cytometry staining and data analysis

The cells were washed by resuspension in 100 μL of 1X DPBS per well to remove the remaining medium. The plate was centrifuged at 300x*g* for 5 min at 4°C, and the supernatant was removed. The Fc receptors on the phagocytes were blocked by resuspending the cells in 25 μL of Fc-blocking antibodies (produced in-house) at a concentration 10 μg/mL, and incubating for at least 10 min on ice. After blocking, 65 μL of staining buffer (1X DPBS supplemented with 2% FCS and 0.005% sodium azide) and 10 μl of BD Horizon™ Brilliant Stain Buffer Plus per well were combined, containing Viakrome 808 (Beckman Coulter, cat. C36628) (1:1000) and a corresponding concentration of fluorophore-conjugated antibodies (Table S1) to the final volume of 100 μl per well. The cells were resuspended in staining buffer, and the plate was incubated for 30 min at 4 °C protected from the light. The identification of rare events can be challenging due to intrinsic cellular autofluorescence, which can vary substantially between the cell types within the same organ (72–75). Autofluorescence is most prominent in macrophages but is also attributed to other myeloid cells (73,76–80). In our experiments on splenic autofluorescent cells, we find that the autofluorescence is attributed to F4/80^+^ RPMs (data not shown), which is in line with the existing literature (78,81). Therefore, we opted for ‘label-free’ identification of RPMs. Next, the plate was centrifuged at 300x for 5 min at 4°C, and the supernatant was removed. The cells were washed in 100 μl of 1X DPBS to remove the remaining medium and centrifuged at 300x*g* for 5 min at 4°C, and the supernatant was removed. The washing step was repeated two more times. Finally, the cells were resuspended in 100 μl of FACS buffer and the plate was stored at 4 °C protected from the light until measurement. From each sample, 85 μl were recorded after obtaining a successful QC (following the manufacturer’s recommendations). Data files were exported and formal analysis was performed in FlowJo Software v.10 (FlowJo Treestar, LLC, Ashland, OR, USA). Minimum Information about a Flow Cytometry Experiment (MIFlowCyt) Checklist is provided in the Supplementary information (Data S1).

## Statistics

For comparing organ distribution based on % dose, the following groups were compared via two-way ANOVA followed by Dunnett’s post hoc test: DSPC:DSPG:CHOL IV 1h vs. DSPC:DSPG:CHOL IV 24h, DSPC:DSPG:CHOL IV 1h vs. DOPC:DOPG:CHOL IV 1h, and DSPC:DSPG:CHOL IV 24h vs. DSPC:DSPG:CHOL SC 24h.

To compare the frequencies of identified subsets as shown on the gating strategy (**Figure S2**) in mice between treatment groups, cell subset frequencies of CD45^+^ cells were exported and analysed using GraphPad Prism 10. All groups were compared via one-way ANOVA followed by Turkey’s post hoc test, except for DOPC:DOPG:CHOL IV 1h vs. DSPC:DSPG:CHOL IV 1h and DOPC:DOPG:CHOL IV 1h vs. DSPC:DSPG:CHOL SC 24h.

To assess Cy5-positive cells per identified cell subset, cells were pre-gated as shown in **Figure S2**. For each end-point cell population, the Cy5-positive gate was set based on a sample from a PBS-injected animal. The frequency of Cy5-positive cells for each identified cell subset from fluorescent liposome-injected mice was exported from FlowJo into an Excel file, and analyzed using GraphPad Prism 10. To compare the frequencies of Cy5-positive cells of each cell subset across experimental groups, a two-way ANOVA followed by Dunnett’s post hoc test (p ≤ 0.05) was performed, using DSPC:DSPG:CHOL IV 1h as the reference group. DOPC:DOPG:CHOL IV and DSPC:DSPG:CHOL IV 24 h were compared individually to the reference group.

To compare the brightness of the liposome signal per cell subset, the (geometric) mean fluorescence intensity (MFI) values of each identified cell subset (pre-gated as shown in **Figure S2**) were exported from FlowJo into an Excel file and analyzed using GraphPad Prism 10. The average MFI per cell subset was calculated for each experimental group of mice. To correct for cell subset-specific autofluorescence, the MFI of each cell subset from the PBS-treated control group (lacking Cy5 fluorescence) was subtracted from the corresponding MFI in the experimental groups. This subtraction yielded the autofluorescence-corrected reporter protein signal per cell subset.

## Results

### Whole organ distribution of PG-containing liposomes

One hour after IV injection of DSPC:DSPG:CHOL and DOPC:DOPG:CHOL liposomes, around 65% and 90% of the liposomes, respectively, remained in the circulation. At this time point, only a small fraction of DOPC:DOPG:CHOL liposomes was detected in the liver (7%) and a very low signal was found in the spleen (1%) (**Table 2**). However, DSPC:DSPG:CHOL liposomes were already present in the liver (22.5% of injected dose) and spleen (11%) at significant doses at this time-point. After 24 hours, the IV injected DSPC:DSPG:CHOL liposomes were not detectable in the blood (**Figure 2**), while 45% of the initial dose was found in the liver and only 2% in the spleen. One-half of the initial dose was not detected in any tested organs. After 24h after SC-injected liposomes, the liposome fluorescence was not detected in the spleen, liver, or blood (**Table 2, Figure 2**). For all of the liposome-injected groups, no liposome fluorescence signal was present in the lungs (**Table 2**). Low average fluorescence with high standard deviation was detected in some lymph nodes and bone marrow; however, the result was not significant (**Table 2, Figure S1**).

**Table 1:** Physicochemical characterization of Cy5-labelled liposomes. All showing mean ± SD, n = 2.

| Formulation | Z-ave (nm) | PDI | Z-potential (mV) |
| --- | --- | --- | --- |
| DOPC:DOPG:CHOL:DSPE-Cy5 | 167.2 ± 3.3 | 0.12 ± 0.04 | -55.4 ± 6.7 |
| DSPC:DSPG:CHOL:DSPE-Cy5 | 194.3 ± 9.3 | 0.07 ± 0.02 | -53.7 ± 10.8 |

**Table 2:** Total organ fluorescence (AU), measured signal corrected for background (PBS-injected mice), n = 3 mice per group.

|  | DSPC:DSPG:CHOL IV<br>1h | DOPC:DOPG:CHOL IV<br>1h | DSPC:DSPG:CHOL IV<br>24h | DSPC:DSPG:CHOL SC<br>24h |
| --- | --- | --- | --- | --- |
| Organ | Total organ fluorescence (AU) ± SD | | | |
| Blood | 3452145 ± 1172763 | 509632 ± 554262 | 0 ± 0 | 0 ± 0 |
| Liver | 1111115 ± 71148 | 376487 ± 186566 | 1704916 ± 63105 | 0 ± 0 |
| Spleen | 540807 ± 189104 | 57950 ± 30090 | 83240 ± 61936 | 11875 ± 16794 |
| bLN | 0 ± 0 | 0 ± 0 | 641 ± 1111 | 0 ± 0 |
| iLN | 1390 ± 314 | 2067 ± 2134 | 905 ± 950 | 1270 ± 608 |
| pLN | 0 ± 0 | 263 ± 228 | 1356 ± 2349 | 1084 ± 1533 |
| mesLN | 67 ± 115 | 0 ± 0 | 0 ± 0 | 346 ± 489 |
| Bone marrow | 18 ± 31 | 4030 ± 3802 | 979 ± 1695 | 1742 ± 2823 |
| Lungs | 0 ± 0 | 0 ± 0 | 0 ± 0 | 0 ± 0 |

**Figure 2:**
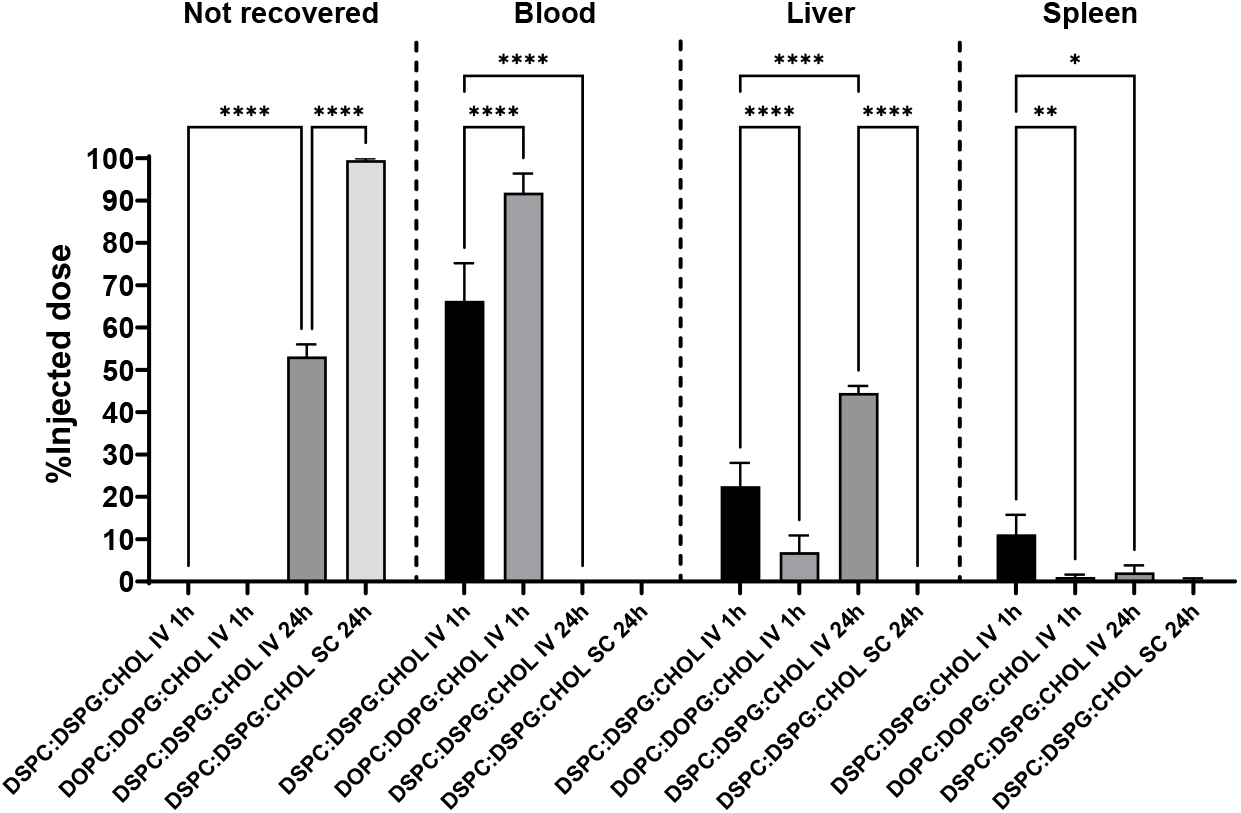
Biodistribution of Cy5-labelled liposomes after IV or SC injection. Only organs with >0.1% dose are shown. N = 3. Means ^+^ SD, ****p < 0.0001, **p < 0.01, *p < 0.05, determined by two-way ANOVA and Tukey’s multiple comparisons test.

### Characterization of splenic immune cells using flow cytometry

While total organ fluorescence is commonly used to assess the distribution of fluorescently labeled drugs across the body, it offers limited information on the specific cell subsets involved in the uptake process (**Figure 1**). To gain a deeper understanding of liposome distribution across cell subsets, a flow cytometry panel was developed to characterize splenic immune cells and detect the association of fluorescently labelled liposomes with delineated cell subsets. Firstly, we selected critical surface markers required for delineation of immune cell subsets of interest (**Table S1**). After marker selection, antibody-fluorophore pairing and titration were performed to select compatible fluorophores and prevent sample overstaining, respectively (**Table S2**). Minimal spillover was allowed into the red detector, reserved for the detection of red-emitting fluorophores, i.e. Cy5, to study the biodistribution of fluorescently labeled liposomes across cell subsets. The representative gating strategy is depicted in **Figure S2**. This gating strategy was applied to identify subsets of macrophages – CD169^+^ marginal zone macrophages, MARCO^+^ marginal zone macrophages, and red pulp macrophages (RPMs), several subsets of DCs – plasmacytoid DCs (pDCs), conventional DCs – DCs type 1 (cDC1s), and cDC2s, and other (non-conventional) DCs, and the additional immune cells such as neutrophils, B cells, monocytes/myeloid cells, and T cells.

Next, we also quantified cell frequencies for each experimental group and compared them among the groups (**Figure S3**). While the identification of the more abundant (>5%) cell subsets (B and T cells, RPMs) was not significant, the frequency of three low-abundance subsets differed among the groups. Specifically, the frequency of MARCO^+^ macrophages, cDC1s, and myeloid cells was significantly different. Mice IV-injected with DSPC:DSPG:CHOL liposomes 24h earlier exhibited significantly higher frequency of MARCO^+^ macrophages compared to the control group and other groups injected with the same liposomes, as well as significantly lower cDC1 cell frequency compared to mice injected with the same liposomes (1h, IV). Lastly, PBS group had a significantly higher frequency of myeloid cells in the spleen than mice IV-injected with DSPC:DSPG:CHOL (**Figure S3**).

### Splenic immune cell distribution of PG-containing liposomes

Liposome distribution was analyzed on a single-cell level to obtain detailed information on which immune cell subsets in the spleen interact with rigid DSPC:DSPG:CHOL (1h and 24h post IV injection) and less rigid DOPC:DOPG:CHOL liposomes (1h post IV injection). Since DSPG liposomes administered SC did not reach the spleen efficiently (**Figure 2, Table 2**), only the results from IV-injected liposome groups are considered. Cells were pre-gated as shown in the gating strategy (**Figure S2**), and liposome-positive gates were set to have <1% of events based on PBS-injected mice.

First, we compared the frequency of Cy5-positive live splenocytes between the groups (**Figure 3**, “Live splenocytes”). Within the first hour after IV injection, 44% of total live splenocytes were associated with DSPC:DSPG:CHOL liposomes, whereas after 24h after injection, the frequency of liposome-associated immune cells drops by almost half (from 44% to 24%). The association is considerably lower with DOPC:DOPG:CHOL liposomes, where only 14% of live splenocytes were detected as positive for Cy5 label within the first hour.

**Figure 3:**
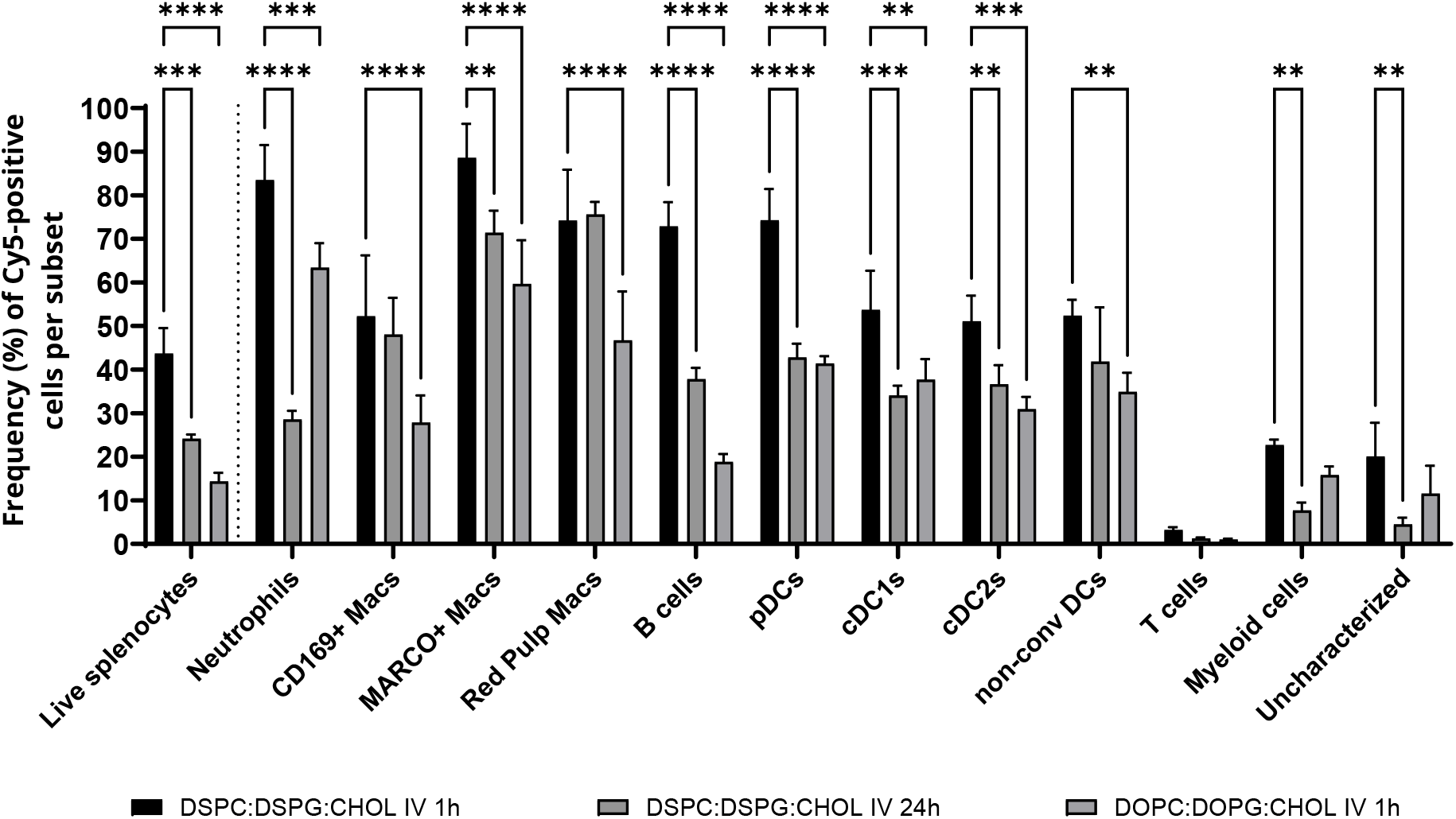
Frequency of Cy5-positive cells among all live splenocytes, or within each cell subset, analysed using flow cytometry, and identified as shown on the gating strategy (Figure S2). Means ^+^ SD, ****p < 0.0001, ***p<0.001, **p < 0.01, *p < 0.05, determined by two-way ANOVA and Dunnett multiple comparisons test. N = 3.

To elucidate the homogeneity of liposome uptake, we compared the frequency of Cy5-positive cells in each cell subset per treatment group (**Figure 3**). Within 1 hour after DSPC:DSPG:CHOL liposome injection, over 70% of neutrophils, MARCO^+^ and RPMs, B cells, and pDCs were Cy5-positive, and around 50% of CD169^+^ macrophages and DCs exhibited Cy5 signal. After 24 hours, the frequency of DSPC:DSPG:CHOL liposome-positive CD169^+^ macrophages, RPMs, and non-conventional DCs remained unchanged, while other cell subsets exhibited a significant decrease in fluorescence, especially neutrophils, B cells, and pDCs (p < 0.0001). Cellular association with less rigid DOPC:DOPG:CHOL liposomes was consistently lower compared to DSPC:DSPG:CHOL liposomes for all identified cell subsets, with the most pronounced decrease in Cy5 fluorescence detected in B cells and pDCs (p < 0.0001), followed by MARCO^+^ and RPMs (p < 0.001). Lastly, neither of the treatment groups showed liposome association with T cells, (**Figure S3**), nor with non-hematopoietic (CD45-negative) splenocytes, which represent <10% of the cells in the spleen (data not shown).

While the percentage of Cy5-positive cells indicates whether a cell has taken up liposomes or not, it does not reflect the level of liposome uptake (binding and internalization) per cell type. Liposome uptake was quantified by Cy5-liposome-specific fluorescence per identified cell subset and compared among the groups. The averaged mean MFI of Cy5 label per cell subset for each experimental group of mice was normalized for cell subset-specific autofluorescence based on the PBS group. In all three investigated treatment groups, splenic macrophages (CD169^+^, MARCO^+^ and RPMs) accounted for over half of the total fluorescence, with MARCO^+^ macrophages exhibiting the brightest fluorescence. In case of DSPC:DSPG:CHOL liposomes, a ∼4-fold drop in total liposome fluorescence is observed between 1-hour and 24-hour time points (**Figure 4**), consistent with a drop of 11% injected dose to 2% injected dose from total spleen fluorescence measurements (**Figure 2**). Changes in relative contribution to total liposome fluorescence were observed for MARCO^+^ macrophages (from 60% to 40%), neutrophils (from 14% to 4%), and pDCs (from 5.4% to 3.8%), while the contribution of RPMs to the total fluorescence increased (from 9% to 30%) (**Figure 4**). We further assessed the retained Cy5 fluorescence in cell subsets that contributed to more than 1% of the total MFI, thereby excluding T cells (0.2%), myeloid cells (0.6%), and uncharacterized cells (0.5%) (**Figure S4A**). RPMs exhibited the highest retention (77% of dose retained at 24 hours compared to 1 hour), followed by non-conventional DCs (56%), cDC2s (54%), CD169^+^ macrophages (48%), cDC1s (38%), B cells (26%), pDCs (17%), MARCO macrophages (16%), and neutrophils (7%). Lastly, we compared the average cell subset abundance with the Cy5 MFI, detected 24 hours after IV injection of DSPC:DSPG:CHOL liposomes (**Figure S4B**). While highly abundant B cells and low-abundance DC subsets exhibited a similar level of Cy5 fluorescence, all three macrophage subsets showed the brightest Cy5 signal.

**Figure 4:**
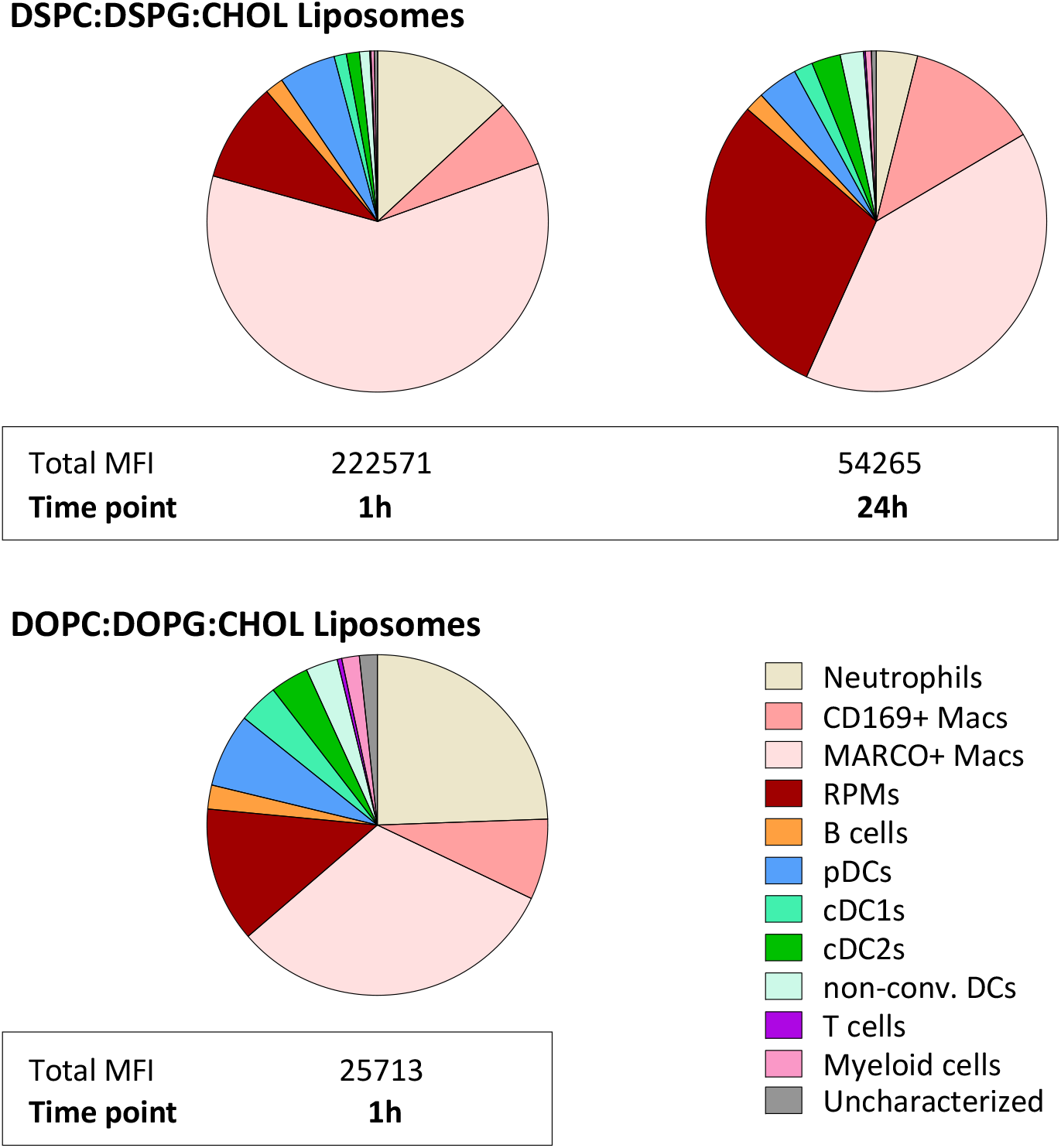
Mean Fluorescence Intensity (MFI, N = 3) of fluorescently-labelled (Cy5) liposomes per cell subset, analyzed using flow cytometry, and identified as shown on the gating strategy (Figure S2).

Lastly, the effect of liposome rigidity on the uptake was evaluated one hour after IV injection. The total mean Cy5 intensity of immune cells associated with rigid DSPC:DSPG:CHOL liposomes was 8.6-fold higher compared to less rigid DOPC:DOPG:CHOL liposomes, in line with organ distribution results (**Figure 2, Table 2**). Besides macrophage subsets, neutrophils exhibited high Cy5 fluorescence in the DOPC:DOPG:CHOL liposome group, contributing to a larger portion of Cy5 fluorescence compared to DSPC:DSPG:CHOL liposomes. Besides neutrophils, the brightest liposome signal was detected in pDCs. Interestingly, B cells, despite accounting for over 50% of liposome-positive cells, exhibited only about 2% of the total Cy5 fluorescence.

## Discussion

Liposomes are a promising drug delivery platform for the treatment of autoimmune and inflammatory diseases (reviewed in (3,82)). We (5) and others (83–86) have reported the efficacy of anionic antigen-carrying liposomal treatment for autoimmune diseases in preclinical models. Still, the exact cellular mechanisms and targets of negatively charged DSPG-containing liposomes remain elusive. In this study, we examined the organ- and splenic cell subset-specific distribution of fluorescently labeled liposomes in Balb/c mice. We addressed several key questions relating to the administration route and liposome rigidity. Primarily, we examined the distribution differences of DSPC:DSPG:CHOL liposomes 1 and 24 hours following IV injection, which allowed us to delineate the temporal dynamics of their distribution. Secondly, knowing that rigid DSPG:DSPC:CHOL liposomes efficiently induce tolerance compared to more flexible DOPC:DOPG:CHOL liposomes (69), we investigated the differences in their distribution 1 hour after IV injection. Lastly, we explored how the administration route (either IV or SC) determines the distribution of DSPC:DSPG:CHOL liposomes 24 hours after injection.

First, we analyzed the fluorescence distribution at the whole-organ level. DSPC:DSPG:CHOL liposomes were rapidly cleared from circulation; after 1 hour, about 65% of the dose remained in circulation, and after 24 hours, none was detected (**Figure 2**). Furthermore, after 24 hours, approximately 50% of the dose was not detected in any of the measured organs, presumably having been eliminated, most likely through excretion pathways. Other studies also observed a rapid clearance of PC:PG:CHOL liposomes from the blood 4 hours after IV injection (87), with almost none detected after 24 hours (88). Interestingly, with increasing time, the liposomes accumulate in the liver, doubling from 23% to 45% of the injected dose. PG liposomes have been described to be taken up almost exclusively by Kupffer cells in the liver, avoiding uptake by hepatocytes (89). At the same time, they are less efficiently retained in the spleen, decreasing from 11% to 2% of the injected dose. Others also observed an accumulation of PC:PG:CHOL liposomes in the liver, hypothesized to be the site of excretion. We could detect no liposome signal in the lungs for any of the groups (**Table 2**). A lack of retention of PC:PG:CHOL liposomes in the lungs, heart, kidneys, and GI tract has been observed before (87,90). Corrected for the weight of the organs, the accumulation of liposomes in the LNs and bone marrow seemed more substantial; however, no differences were found between the groups (**Figure S1**). At this scale, the difference in uptake of DSPC:DSPG:CHOL liposomes in the spleen over time was the most striking, and we further assessed the liposome dynamics in the spleen on the single-cell level.

Using a specially designed and optimized flow cytometry panel, we identified average cell subset frequencies comparable to those reported before (43,44,46–50,52) with some exceptions (**Figure S3**). The frequency of MARCO^+^ macrophages was significantly higher in mice 24h after IV-injection of DSPC:DSPG:CHOL liposomes (**Figure S3**). Early studies identified MARCO as the main receptor involved in the uptake of silica particles in alveolar macrophages in mice (91), and other studies indicate that MARCO is upregulated in tolerogenic APCs. Specifically, research shows that the MARCO gene is upregulated in tolerant macrophages compared to naïve macrophages upon LPS (92,93) and LTA stimulation (93), and partially contributes to the increased phagocytosis (93). MARCO gene expression was also found to be upregulated in DCs pulsed with dead cells (94). Furthermore, a recent study showed that cancer cells induce MARCO expression in tumor-associated macrophages, leading to regulatory/tolerogenic macrophage polarization (95). Future research should therefore examine liposome-dependent upregulation of MARCO and consequent (tolerogenic) effects. Next, we detected a significant decrease in the frequency of other myeloid cells in mice IV-injected with DSPC:DSPG:CHOL liposomes (both 1h- and 24h-time points) compared to PBS mice. This population, mostly consisting of monocyte/macrophage cells (48,96), exhibited low association with liposomes (**Figure 3-4**). The changes in its frequency possibly suggest liposome-specific effects, such as migration (97) of CD11b^+^ cells from the spleen upon liposome encounter, however, this requires further study (**Figure S3**). Furthermore, we observed a lower frequency of cDC1 cells in mice injected with DSPC:DSPG:CHOL liposomes 24h earlier compared to mice from other groups, however, the decrease was significant only in comparison with mice injected with the same liposomes (1h IV) (**Figure S3**). While cDC1s associated DSPC:DSPG:CHOL liposomes (**Figure 3**), they did not exhibit high liposome signal (**Figure 4**) or retainment (**Figure S4**). Still, cDC1s are a migratory population expressing an integrin CD103, which could explain the migration out of the spleen, and can generate regulatory T cells (98,99), indicating their possible involvement in the tolerance induced with DSPC:DSPG:CHOL liposomes.

Using flow cytometry to detect liposomal Cy5 fluorescence, we obtained two types of information: which cell subsets associate with liposomes (liposome-positive cells), and the relative amount of liposomes per cell) (**Figure 2A**). We detected a 1.8-fold decrease in DSPC:DSPG:CHOL liposome-associating splenocytes within 24 hours after injection (**Figure 3**), indicating that liposomes interact with the cells but are either not internalized or are cleared within 24h. The frequency of most liposome-associating cell subsets decreased significantly within 24 hours, except for CD169^+^ macrophages, RPMs, and other non-conventional DCs (**Figure 3**). These subsets appear to associate with and retain liposomes over time, indicating their functional involvement in liposome-induced effects.

We further quantified the liposome association efficiency by assessing the fluorescence intensity of the liposome label per single cell. All identified macrophage subsets, RPMs, MARCO^+^, and CD169^+^ macrophages, contributed to the highest proportion of liposome fluorescence in the spleen 1 h and 24 h after injection, confirming their superior phagocytic capacity compared to other APCs (**Figure 4**) (101,102). Cells exhibiting the highest retention of liposome fluorescence between the two time points were RPMs (76.5%), non-conventional DCs (55.5%), cDC2s (53.9%), and CD169+ macrophages (47.9%) **(Figure S4A**). While this result indicates that the identified cell subsets are primarily involved in the initial association and uptake of IV-injected DSPC:DSPG:CHOL liposomes, we cannot confirm initial capture and retention of liposomes over time using the presented technique. Further testing should be performed to elucidate whether the same effects apply to other types of liposomes or nanoparticles.

Considering the frequency of splenic cell subsets and their relative contribution to the total Cy5 fluorescence, macrophage subsets, especially MARCO^+^ macrophages, exhibit high liposome uptake 24 hours after liposome administration (**Figure S4**). Similar findings on the association of PS-containing liposomes with MARCO^+^ and CD169^+^ macrophages at the same time-point have been reported using microscopy (52). Other cell subsets also associate with liposomes, albeit less efficiently (**Figure 4**). Neutrophils contribute to 13% of the total liposome signal within the first hour, but retain only 7% of this fluorescence after 24 hours (**Figure S4A**). B cells, the most abundant cell population in the spleen, only contribute to a small proportion of total Cy5 fluorescence (**Figures 4** and **S4B**). DCs also contributed to a relatively low proportion of liposome fluorescence. While pDCs exhibited the brightest liposome fluorescence among the DC subsets, followed by cDC2s at the 24h time point (**Figure 4**), cDC2s and non-conventional DCs showed higher liposome retention (**Figure S4A**). Splenic cDC2s reside in the MZ exposed to the blood flow (103) where they can take up liposomes, and consequently migrate to T cell zones to initiate T cell responses (104,105). Therefore, we cannot exclude the important functional involvement of DCs in the liposome uptake process of rigid DSPC:DSPG:CHOL liposomes.

We further explored the impact of liposome rigidity on uptake at the whole-organ and single-cell level. DOPC:DOPG:CHOL liposomes circulated to a much higher extent than the more rigid DSPC:DSPG:CHOL liposomes, with around 90% of the dose still detectable in the blood after 1 hour (**Figure 2**). DOPC:DOPG:CHOL liposomes have lower uptake in APCs than DSPC:DSPG:CHOL liposomes (69), and may therefore be cleared less rapidly. Anselmo *et al*. also initially observed a higher clearance of hard hydrogel nanoparticles compared to soft particles, although no differences were observed after 4 hours (106). Since almost all of the DOPC:DOPG:CHOL liposomes were still circulating after 1 hour, it is not surprising to find only a small proportion of the injected dose in the liver (7%) and spleen (1%). Despite a dim fluorescence detected in the spleen by a fluorescent plate reader (**Table 2**), a bright fluorescence signal was detected using flow cytometry, allowing us to assess liposome association on a single-cell level. We observed over 8-fold lower accumulation of unsaturated liposomes compared to saturated liposomes in the spleen within 1 hour. The most striking difference in Cy5 fluorescence distribution was that a higher proportion of DOPC:DOPG:CHOL liposomes was associated with neutrophils and a lower proportion with MARCO^+^ macrophages compared to DSPC:DSPG:CHOL liposomes.

Comparing injection routes, the main observation is that almost none of the dose was recovered after SC injection (99.6% not detected, **Figure 2**), and liposome fluorescence in the spleen was not detectable using flow cytometry (data not shown). This is most likely due to the liposomes being retained at the site of injection 24 hours after injection. 52 hours after SC injection in the flank (the same site as used in this study), around 95% of the injected EPC:EPG:CHOL liposomes dose was still at the site of injection (12), and almost none of the EPC:EPG:CHOL liposome dose was found in the liver, spleen, and LNs (12,107), which aligns with our data.

## Conclusion

Overall, we have developed a flow cytometry panel to assess in detail the uptake of Cy5-labelled nanoparticles in rare APC subsets in murine splenocytes. The application of this technique provides a deeper understanding of liposome biodistribution and uptake by splenic cells, suggesting that future research should focus on the role of macrophages in liposome-induced immune tolerance and further investigate the impact of liposome characteristics, such as bilayer rigidity and composition on their therapeutic potential.

## Author contribution

Deja Porenta: conceptualization, formal analysis, data curation, writing-original draft. Estefanía Lozano-Andrés: methodology, writing-review, and editing. Enrico Mastrobattista: writing-review and editing. Femke Broere: writing-review and editing. Naomi Benne: conceptualization, methodology, supervision, writing-review and editing.

## Data availability

Minimum Information about a Flow Cytometry Experiment (MIFlowCyt) Checklist is provided in the Supplementary information (Data S1). Flow cytometry files (.FCS) are available upon request to the corresponding author.

## Acknowledgments

The authors thank the Flow Cytometry and Cell Sorting Facility at the Faculty of Veterinary Medicine at Utrecht University for support. E.L.A. is an International Society for Advancement of Cytometry (ISAC) SRL Emerging Leader 2025-2028.

