## Supplements for "Detailed splenic single-cell biodistribution of phosphatidylglycerol-containing liposomes"

**Supplementary materials**

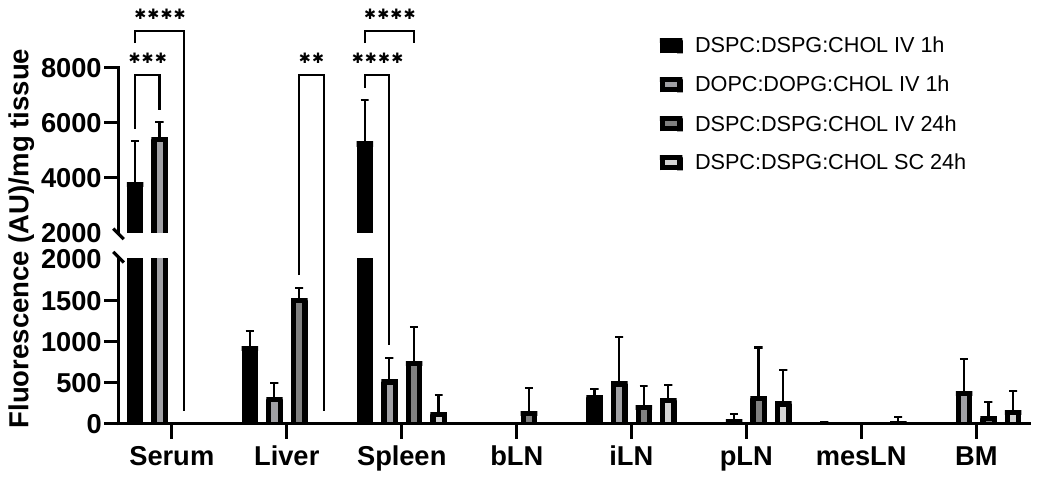

**Figure S1: Fluorescence of Cy5-labelled liposomes after IV or SC injection corrected for organ weight.** Only organs with >0 AU fluorescence are shown. N = 3. Means ^+^ SD, ****p < 0.0001, ***p < 0.001, **p <0.01 determined by two-way ANOVA and Tukey’s multiple comparisons test.

**Table S1: Reagents and antibodies used for flow cytometry staining**

| **Subset** | **Phenotype*** |
| --- | --- |
| Neutrophils | Ly6G^+^ SSC^-^A^intermediate^ |
| Eosinophils | SSC^-^A^high^ |
| Marginal metalophilic macrophages | SSC^-^A^low^ CD169^+^ |
| Marginal zone macrophages | SSC^-^A^low^ CD169^-^ MARCO^+^ |
| Red pulp macrophages (RPMs) | SSC^-^A^low^ CD169^-^ MARCO^-^ Autofluorescence^high^ |
| B cells | SSC^-^A^low^ CD169^-^ MARCO^-^ CD11c^-^ B220^+^ |
| plasmacytoid DCs (pDCs) | SSC^-^A^low^ CD169^-^ MARCO^-^ B220^+^ CD11c^+^ |
| conventional DCs type 1 (cDC1s) | SSC^-^A^low^ CD169^-^ MARCO^-^ B220^+^ CD11c^+^ CD11b^-^ XCR1^+^ |
| conventional DCs type 2 (cDC2s) | SSC^-^A^low^ CD169^-^ MARCO^-^ B220^+^ CD11c^+^ XCR1^-^ CD11b^+^ |
| Non-conventional DCs | SSC^-^A^low^ CD169^-^ MARCO^-^ B220^+^ CD11c^+^ XCR1^-^ CD11b^-^ |
| T cells | SSC^-^A^low^ CD169^-^ MARCO^-^ B220^+^ CD11c^-^ CD11b^-^ CD3e^+^ |
| Monocytes and other myeloid cells | SSC^-^A^low^ CD169^-^ MARCO^-^ B220^+^ CD11c^-^ CD3e^+^ CD11b^+^ |

*Cells were pre-gated on splenocytes, single cells, live cells, and CD45^+^ cells.

**Table S2: Cell subset characterization**

| **Specificity** | **Clone** | **Fluorophore** | **Dilution** | Company | **Cat. number** |
| --- | --- | --- | --- | --- | --- |
| CD45 | 30-F11 | Alexa Fluor 488 | 1:1600 | Biolegend | 103121 |
| CD45R (B220) | RA3-6B2 | Alexa Fluor 700 | 1:400 | Biolegend | 103231 |
| CD3e | 145-2C11 | Brilliant Violet 510 | 1:100 | Biolegend | 100353 |
| Ly-6G | 1A8 | Brilliant Violet 510 | 1:300 | Biolegend | 127633 |
| CD11b | M1/70 | eFluor 450 | 1:800 | Invitrogen | 14-0112-82 |
| CD11c | N418 | PE-Vio770 | 1:200 | Miltenyi Biotec | 130-120-297 |
| XCR1 | ZET | Brilliant Violet 785 | 1:200 | Biolegend | 148225 |
| CD169 | 3D6.112 | BV605 | 1:200 | Biolegend | 142413 |
| MARCO | # 2359A | Alexa Fluor 594 | 1:200 | R&D Systems | FAB29561T |
| Viability | n/a | Viakrome 808 | 1:1000 | Beckman Coulter | C36628 |

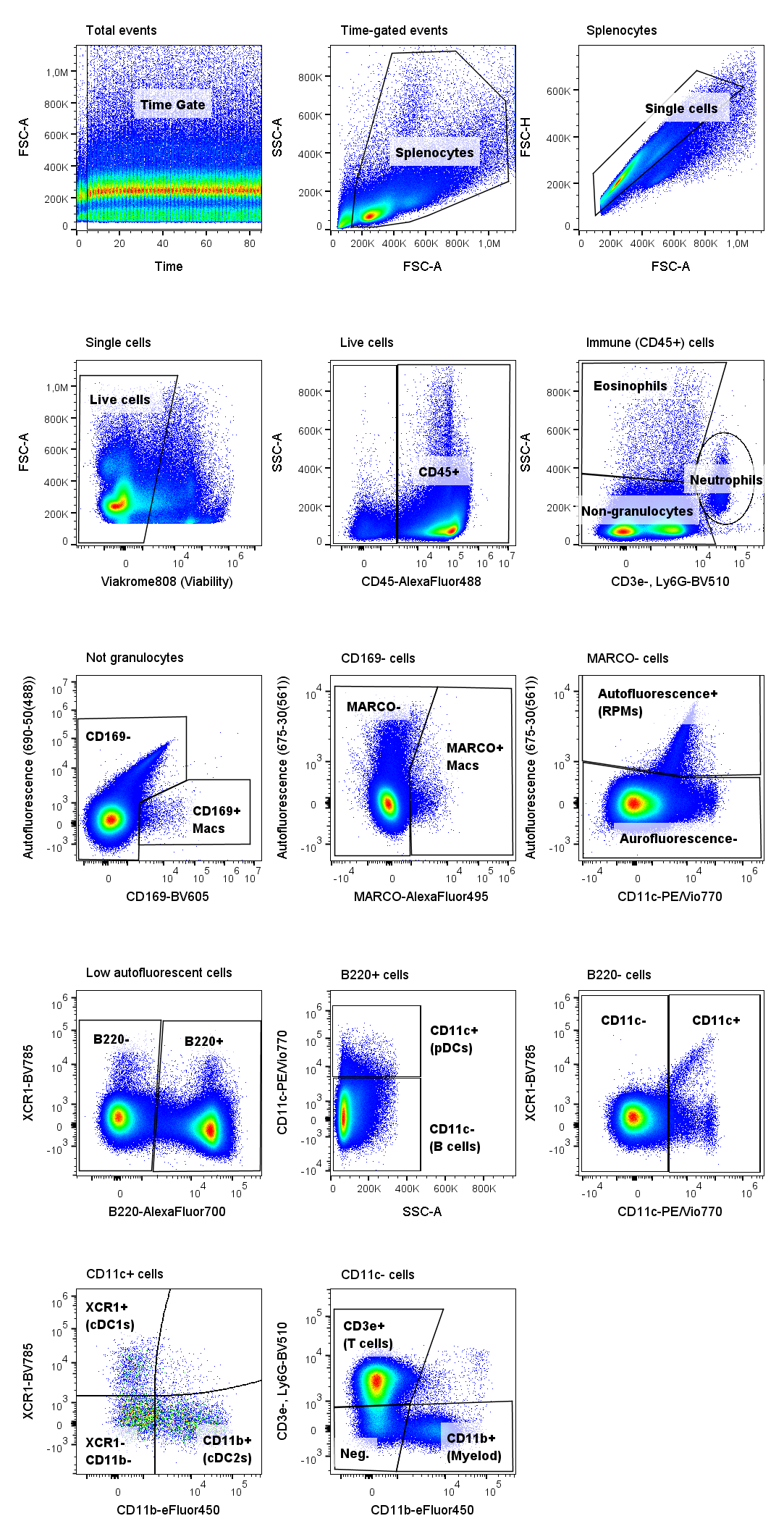

**Figure S2: Gating strategy for characterization of splenic immune cells in healthy mice.** Spleens were mechanically processed through a cell strainer to generate single-cell suspensions. Cells were treated with Fc-blocking antibodies, followed by surface staining with fluorophore-conjugated antibodies. Cells were pre-gated as shown: time, splenocytes, single, live cells. From this point forward, the displayed plots represent concatenated samples (n=3). Hematopoietic (immune) cells are identified by CD45 staining. Neutrophils are identified among CD45^+^ cells as Ly6G^+^ with intermediate side scattering, and eosinophils as SSC-A^high^. CD169^+^ macrophages are identified among ‘Not granulocytes’, and MARCO^+^ marginal zone macrophages are identified among CD169^-^ cells. Autofluorescent red pulp macrophages (RPMs) are identified based on autofluorescence recorded on B690 detector. B220-positive and -negative cells are identified among ‘Low autofluroescent cells’. Plasmacytoid DCs are identified among B220^+^ cells by the expression of CD11c, while CD11c^-^ cells are characterized as B cells. CD11c^+^ DC-like cells are identified among B220-negative cells, and the population is further plotted as XCR1 vs. CD11b to distinguish conventional DCs type 1 (XCR1^+^ CD11c^+^) and conventional DCs type 2 (CD11b^+^ CD11c^+^) from other, non-conventional DCs (CD11c^+^). Not DCs (CD11c^-^) are plotted across myeloid marker CD11b, to distinguish monocyte/myeloid cell populations, and CD3e, to identify T cells. The remaining cells remain uncharacterized (<4% of CD45^+^ splenocytes). The panel has additionally been tested with Ly-6C (APC-Cy7, dilution 1:1600, Biolegend, #128025), which can be added if desired.

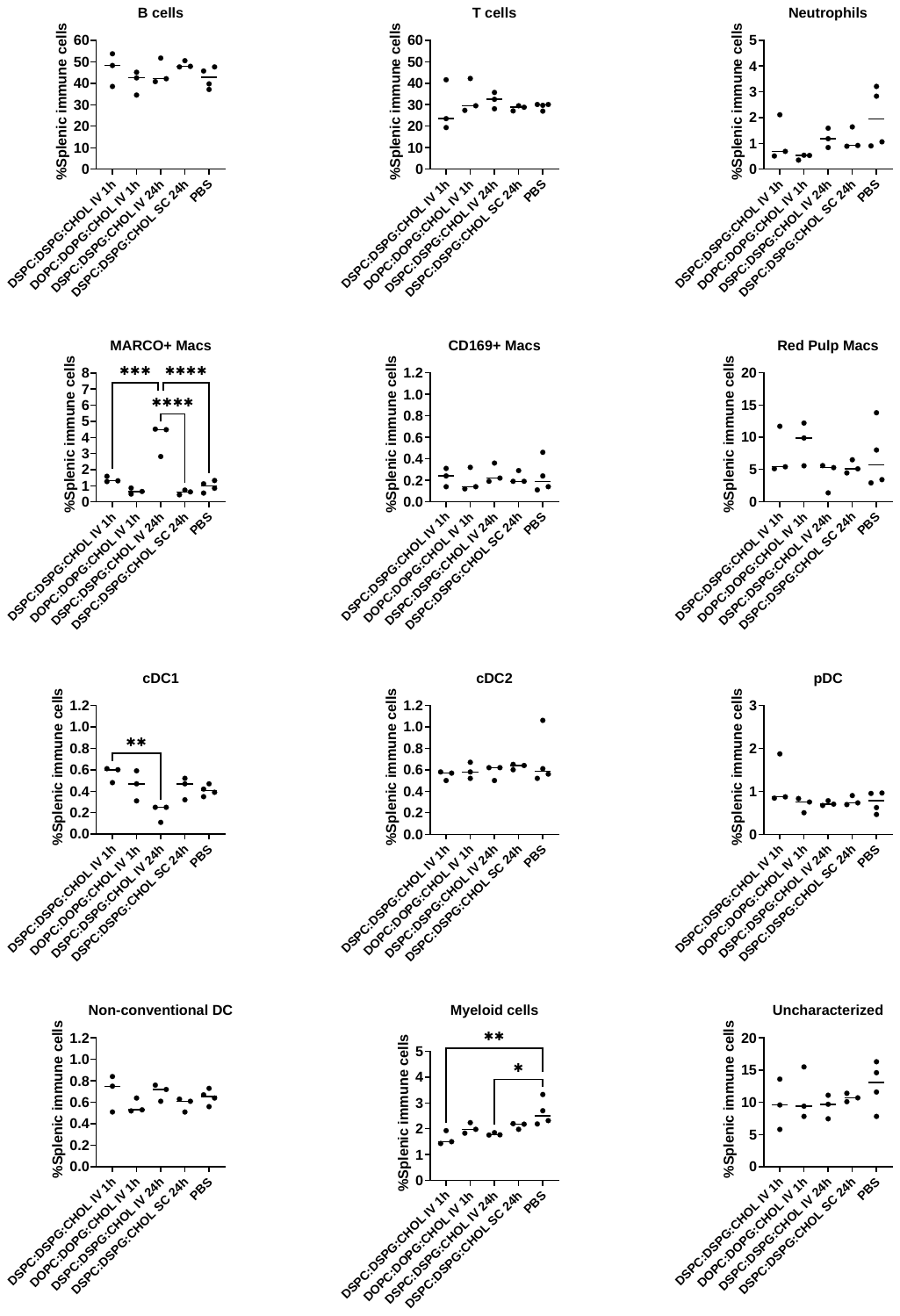

**Figure S3: Frequency of cell subsets among splenic immune cells per experimental group**. Cell subsets were identified as shown on the gating strategy (Figure S2), and the frequencies of CD45^+^ cells were exported and analysed using GraphPad Prism 10. Significance was determined by one-way ANOVA followed by Turkey’s post hoc test (p ≤ 0.05). Experimental groups: N = 3, PBS group: N=4. Means ^+^ SD, ****p < 0.0001, ***p < 0.001, **p < 0.01, *p < 0.05.

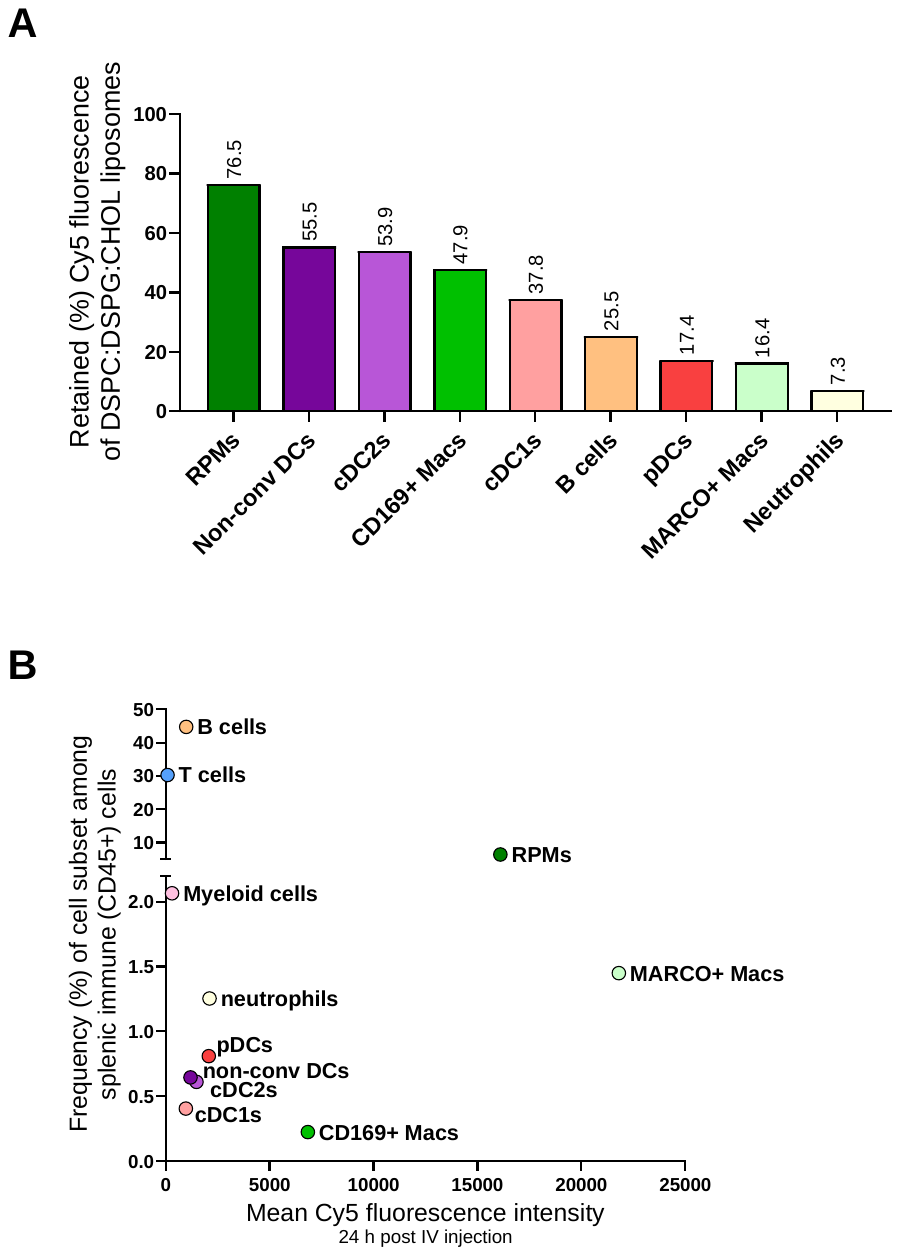

**Figure S4:** **DSPG:DSPG:CHOL liposome retention and fluorescence intensity considering cell subset frequency**. (A) Percentage of retained Cy5 fluorescence of liposomes 24 h after IV injection, relative to the fluorescence measured 1h after injection. Percentages of retained liposome fluorescence 24 h after injection were calculated by considering that the liposome fluorescence measured 1h after liposome administration per cell subset represents 100%. Only cell subsets that contributed to more than 1% of the total MFI are represented, thus excluding T cells, myeloid cells, and uncharacterized cells. (B) Representation of the mean Cy5 fluorescence intensity 24 h after IV injection and the frequency of the corresponding cell subset among all splenic immune cells. The frequency of each cell subset at the 24 h time point was exported and plotted against the mean fluorescence intensity measured per cell subset.
